# A secreted helminth microRNA suppresses gastrointestinal cell differentiation required for innate immunity

**DOI:** 10.1101/2024.12.20.629722

**Authors:** Matias G. Perez, Victoria Gillan, William M. Anderson, François Gerbe, Fabien Herbert, Tom N. McNeilly, Rick M. Maizels, Philippe Jay, Eileen Devaney, Collette Britton

**Author notes:** Corresponding authors: Correspondence addresses: Lead contact.

## Abstract

Pathogens have developed multiple strategies to modulate host immune defense mechanisms. Understanding how this is achieved has potential to inform novel therapeutics for diseases caused by immune dysfunction. Parasitic helminths are masters of immune evasion, via release of secreted products, resulting in chronic infection. Helminths secrete small regulatory microRNA (miRNAs), which can interact with host cells. Here we show that a single parasite miRNA (miR-5352), conserved across gastrointestinal (GI) nematodes, suppresses IL-13-induced GI epithelial cell differentiation and cytokine responses, and promotes stem cell maintenance. Mechanistically, this is achieved through targeted repression of critical host factors, including Klf-4 and the IL-22 receptor, together with modulation of Wnt and Notch signalling pathways. Nematode miR-5352 shows seed sequence conservation with mammalian miR-92a family members, indicating that through convergent evolution, GI nematodes exploit a host miRNA regulatory network to suppress host innate responses, promote tissue regeneration and establish a favourable environment for chronic infection.

## Introduction

To successfully establish and survive for a protracted time in a host, parasites extensively modulate immune defense mechanisms. Helminth parasites that inhabit the gastrointestinal (GI) tract are among the most ubiquitous and successful pathogens. They infect over 1.5 billion people^1^ and are a major cause of livestock production losses globally^2,3^. The success of these parasites can be attributed to their ability to manipulate host immune responses by modifying surrounding tissue and limiting inflammation, leading to a chronic infection with minimal tissue damage^4^. Characterisation of proteins produced and secreted by helminths has led to the discovery of several immunomodulators^5,6^. Little is known, however, about the targets and effects of helminth secreted small RNAs, including microRNAs (miRNAs).

MiRNAs were first identified as regulators of developmental timing in the free-living model nematode *Caenorhobditis elegans*^7,8^. This seminal work defined a fundamental principle governing how gene expression is regulated^9^ (Nobel Prize in Physiology or Medicine 2024^10^). MiRNAs modulate gene expression post-transcriptionally by binding to the 3’UTR of their target mRNAs, resulting in inhibition of protein translation and transcript degradation^11^. The repertoire of miRNAs expressed and secreted by various helminth species have been described ^12,13,14^, stimulating interest that they function as potential regulators of host gene expression. MiRNAs in helminth excretory-secretory (ES) products are present within extracellular vesicles (EVs) or in non-vesicular forms. While helminth EVs can modulate host cell gene expression,^12,15,16,17,18^, the targets and effects of specific secreted helminth miRNAs on host GI cells have not been examined.

Following GI helminth infection, epithelial tuft cells, the first sensors and responders to GI parasites, secrete the cytokine IL-25, which activates type 2 innate lymphoid cells (ILC2s) to release the type-2 cytokines, IL-4 and IL-13^19,20,21^. The resulting tuft and goblet (mucous) cell hyperplasia initiates the weep and sweep response that promotes helminth expulsion^19^. While parasite presence can stimulate epithelial cell responses, some parasites also modulate this response, leading to chronic infection. Nematode ES products are therefore complex, comprising activators and suppressors of host immunity. Secreted products from adult worms of the murine nematode *Heligmosomoides polygyrus* can suppress tuft and goblet cell expansion in response to treatment with IL-4 and IL-13, in vitro and in vivo^22^, but the specific parasite molecules involved have not been identified.

To establish in the gut, parasitic helminths modulate the epithelium of the GI tract. The GI epithelium is ordered into stem cells (SC), transient-amplifying (TA) cells and differentiated cell compartments^23,24^. The unique ability of GI epithelial tissue to continuously regenerate is mediated by SC located in gastric glands or intestinal crypts^25,26^. In the SI, epithelial cell fate depends on the transcription factors Atoh1 and Hes1, which regulate Lgr5^+^ SC to follow secretory or absorptive cell lineages, respectively^27^. Secretory cells include goblet, tuft, enteroendocrine, parietal, surface mucous and Paneth cells, while absorptive cells comprise enterocytes^28^. Additionally, signalling pathways, including Wnt, Notch, Hippo and BMP regulate cell renewal and maintain homeostasis of the GI tract^24,29^. The different cell types help maintain GI epithelial barrier integrity and immune tolerance at steady state, and play specific roles in response to bacterial, parasitic and viral infections^30^. Perturbations in GI epithelial cell activity and tolerance can lead to inflammatory bowel diseases (IBD), such as Crohn’s and ulcerative colitis^31^.

The parasite molecules and mechanisms involved in altering epithelial cell fate are not known. Interaction of parasite miRNAs with host target genes is a mechanism that requires further investigation. We previously reported the miRNA repertoire of the important veterinary nematode *Haemonchus contortus*^32^, a blood-feeding ovine abomasum (gastric) parasite related to human hookworms. *H. contortus* is studied extensively as a model nematode for drug discovery, vaccine development and anthelmintic resistance because of the high-quality genome resources available^33^. We identified a group of 23 miRNAs in ES products released from *H. contortus* parasitic L4 larvae and adult stages, including a cluster containing hco-miR-5352. Intriguingly, this miR-5352-containing cluster is conserved in other GI parasitic nematodes, including human-infective hookworms, but is not reported in tissue dwelling nematodes^34^. Importantly, miRNAs of this cluster could be detected in abomasal tissue and draining lymph nodes of *H. contortus*-infected animals, indicating their release in vivo and potential to modulate host gene expression^34^.

Here, we demonstrate that *H. contortus* ES products and miR-5352 alone can suppress the differentiation of GI epithelial secretory cells required for innate immunity. Transfection of murine and ovine GI organoids with a mimic of miR-5352 maintained organoid stemness and suppressed the effects of IL-13 on tuft and mucous cell differentiation. Mechanistically, this is achieved by repression of key host target genes including transcription factor Krüppel-like factor 4 (*Klf4*), IL-22 receptor subunit alpha 1 (*Il22ra1*) and modulators of Wnt and Notch signalling pathways. Notably, nematode miR-5352 shows seed sequence conservation with mammalian miR-92a family members. Our data indicate that through convergent evolution, GI nematodes hijack a host miRNA regulatory network to suppress host innate responses and promote GI epithelial regeneration and homeostasis, creating an environmental niche for their survival.

## Results

### 1. *H. contortus* ES suppresses IL-13 mediated GI secretory cell differentiation

To determine whether *H. contortus* secreted products can modulate host epithelial cell responses driven by type-2 cytokines, murine and ovine GI organoids were treated with adult worm ES together with IL-13. Ovine organoids were derived from stem cells in the abomasum (gastric tissue), the location in which *H. contortus* parasites reside. After exposure to IL-13 alone, there was a characteristic “budding” phenotype, first observed 12 h after IL-13 exposure and not observed in untreated organoids (**Figure 1A**). IL-13 treatment increased expression of *Pou2f3*, a marker of murine and ovine tuft cells^19,35^, and *Muc6*, a mucin expressed in gastric mucosa^36^ (**Figure 1B**), indicating that IL-13 promotes differentiation of ovine abomasal secretory cells. Expression of *Hes1*, which promotes non-secretory enterocytes and is a downstream target gene of the Notch1 signalling pathway^37^ also increased with IL-13 treatment (**Figure 1B**). In the presence of IL-13 and *H. contortus* total ES, organoids showed significantly reduced expression of *Pou2f3* and *Muc6*, but increased expression of *Hes1*, as observed in murine SI organoids treated with IL-13 plus *H. polygyrus* ES^22^. While *H. contortus* ES alone did not alter *Pou2f3* or *Muc6* expression relative to untreated organoids, expression of *Hes1* was increased (**Figure 1B**). Previous studies reported increased expression of *Hes1* during chronic, but not acute, infection with the murine colonic nematode *Trichuris muris*, associated with an increase in cell proliferation and crypt size^38^.

**Figure 1.**
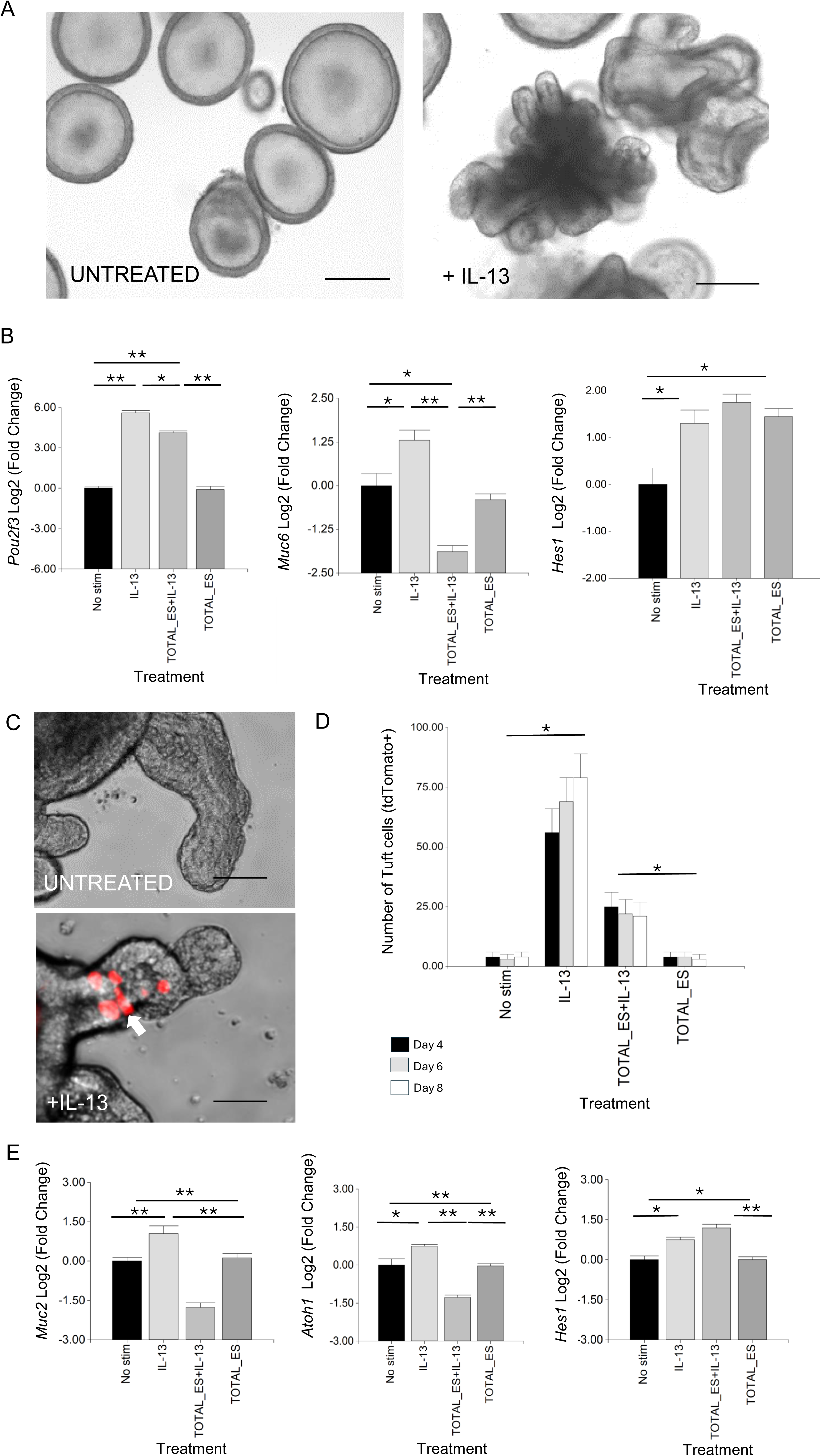
*H. contortus* ES suppresses effects of IL-13 on GI organoid cell differentiation. **(A)** Representative image of ovine abomasal organoids treated or not with IL-13 and cultured in organoid growth medium (OGM) for 4 days. **(B)** RT-qPCR of canonical tuft cell marker *Pou2f3*, mucous cell marker *Muc6* and transcription factor *Hes1* in ovine abomasal organoids. RNA was from unstimulated organoids (No stim) or treated with IL-13 alone, IL-13 plus total *H. contortus* ES or total *H. contortus* ES alone. Log2 fold change of RT-qPCR values compared with unstimulated control in three independent biological replicates +/−SD. **(C)** Representative images of *Dclk1* tdTomato^+^ tuft cells (arrow) in murine SI organoids, untreated or treated with IL-13 in OGM for 4 days at 37°C. **(D)** Number of *Dclk1* tdTomato^+^ cells in murine SI organoids at 4, 6, and 8 days after no stimulation, treatment with IL-13, IL-13 plus *H. contortus* total ES or *H. contortus* total ES alone +/− SD. Graph shows the average number of tdTomato^+^ cells from at least 5 images and in total represents 500 organoids per treatment. Statistics calculated with a two-tailed t-test on mean of biological replicates (n=3) compared to untreated organoids. **(E)** RT-qPCR expression of *Muc2, Atoh1* and *Hes1* in murine SI reporter organoids. RNA was from unstimulated organoids (No stim), those treated with IL-13 alone, IL-13 plus total *H. contortus* ES, or total *H. contortus* ES alone. Log2 fold change of RT-qPCR values are compared with unstimulated control in three independent biological replicates +/− SD. Statistical analysis was by ordinary one-way ANOVA with Tukey’s multiple comparisons test. *H. contortus* total ES at 2 µg/ml. *, P < 0.05; **, P < 0.01. Scale bars = 200 µm.

Effects of *H. contortus* ES on GI epithelial responses to IL-13 were also examined in murine organoids. For this, we employed murine SI organoids in which tdTomato was expressed under the control of the promoter of the tuft cell-associated gene *Dclk1*^19^ (*Ai14; Dclk1-CreERT2*). This facilitated real time imaging of murine SI tuft cells. Following IL-13 treatment, there was a significant increase in tdTomato^+^ cells (10-fold at day 4), which was suppressed by co-incubation with *H. contortus* total ES (**Figures 1C, D**). *H. contortus* ES alone had no effect on the number of SI tuft cells (tdTomato^+^), as observed for ovine abomasal tuft cells. IL-13 treatment also resulted in increased expression of murine SI goblet cell marker *Muc2*^39^, secretory cell promoting transcription factor *Atoh1*/*Math1*^40^ and *Hes1*, which in the SI promotes differentiation of absorptive cells^27^ (**Figure 1E**). These IL-13 mediated effects were modulated by incubation of organoids with *H. contortus* total ES. *Muc2* and *Atoh1* expression were reduced, while *Hes1* levels increased (**Figure 1E**). In the SI, Hes1 negatively regulates Atoh1, resulting in an increase in absorptive cells and reduction in cells of the secretory lineage^40^. *Atoh1* expression was not detected by RT-qPCR in ovine abomasal organoids, suggesting that other transcription factors regulate the epithelial secretory lineage in gastric tissue^41^. Our data indicate that *H. contortus* total ES suppresses IL-13 induced secretory cell development in both murine and ovine GI organoids.

Previous studies reported development of “spheroid” SI organoids which failed to bud or differentiate, in the presence of *H. polygyrus* ES prepared by concentration over a 3kDa cut-off membrane^22,42^. Similar foetal-like undifferentiated regions were also observed in murine SI tissue following *H. polygyrus* infection^43^. We examined the effect of concentrated *H. contortus* ES on ovine abomasal organoids, and on ovine and murine SI organoids, and compared this to treatment with *H. polygyrus* concentrated ES. Spheroids were induced in all three organoid types with ES from either species, albeit less obviously in abomasal organoids which have a low level of budding under standard (control) conditions (**Supplementary Figure S1** at 12 h after incubation with ES). Notably, in the presence of concentrated ES, ovine SI organoids showed greater budding and fewer spheroids compared to murine SI organoids, and there was less spheroid formation in ovine or murine SI organoids treated with *H. contortus* ES compared with *H. polygyrus* ES (**Supplementary Figure S1**). Therefore, at high concentration, ES components from the two nematode species induce spheroid formation, with *H. polygyrus* ES having a slightly more potent effect.

### 2. miR-5352 mimic modulates epithelial cell gene expression and reduces organoid budding

MicroRNA miR-5352 is expressed uniquely in parasitic stages of nematodes infecting the GI tract and of the lumen dwelling lungworm *Dicytocaulus viviparus,* and is present in *H. contortus* and *H. polygyrus* ES ^12,34^. We speculated that this miRNA may modulate host gene expression at mucosal sites, and could be involved in altering organoid responses to IL-13. To test this, we firstly optimised delivery of a synthetic miRNA into GI organoids using a Cy3-labeled mimic of miR-5352 (100 nM, details in Materials & Methods). Transfection with labelled miRNA mimic alone or with lipofectamine resulted in uptake in only a few organoid cells (**Figure 2A**), while inclusion of DharmaFECT reagent increased transfection efficacy, with Cy3 observed in many organoid cells (**Figure 2A**).

**Figure 2.**
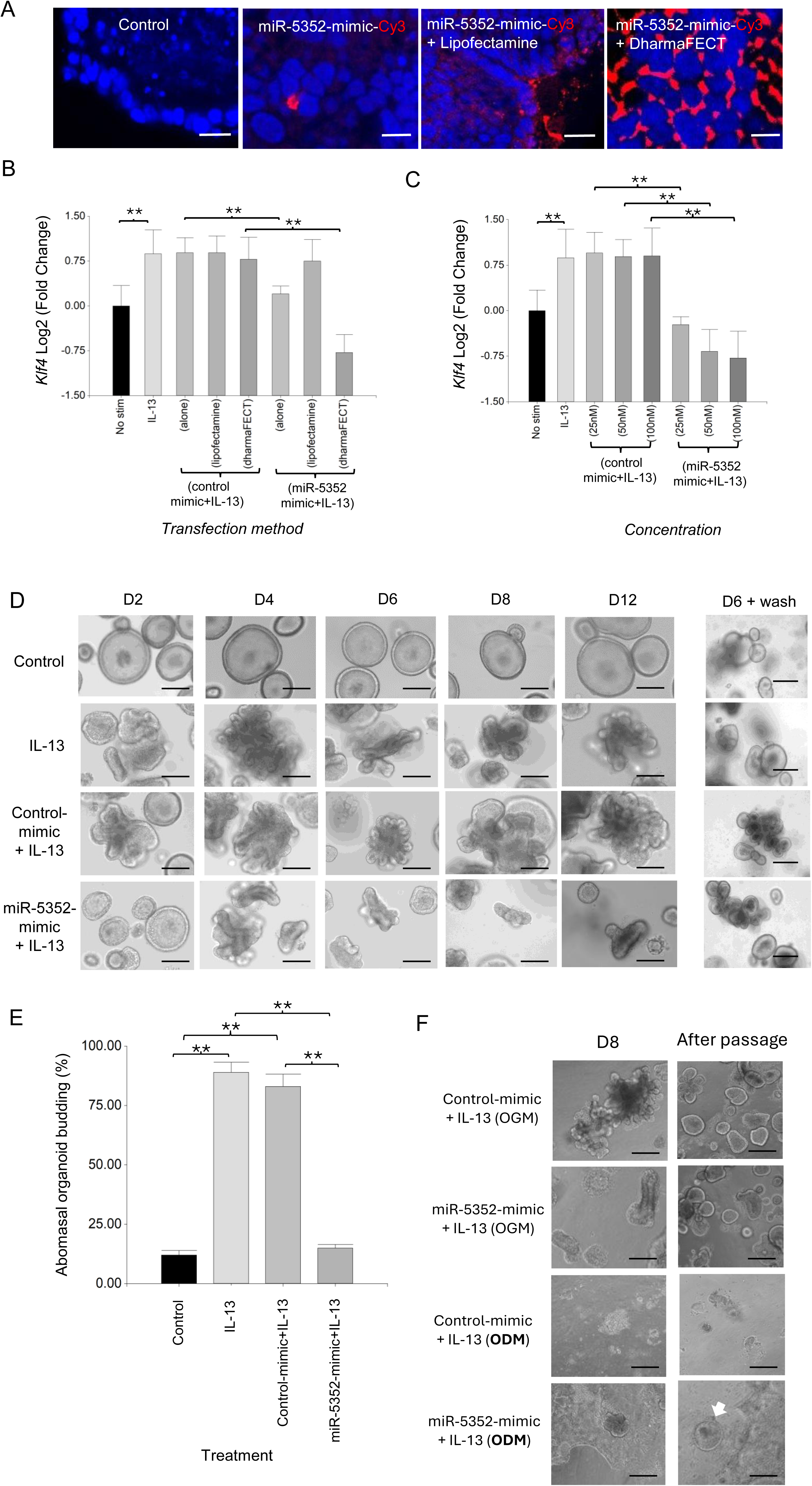
Transfection of ovine abomasal organoids with miR-5352 mimic suppresses IL-13-mediated differentiation. **(A)** Confocal microscopy of DAPI stained ovine abomasal organoids. Control (no treatment) or incubated with miR-5352 mimic-Cy3 alone, miR-5352 mimic-Cy3 + Lipofectamine, miR-5352 mimic-Cy3 + DharmaFECT. Red, Cy3 mimic and blue, DAPI stained nuclei. Scale bars = 10 µm. **(B)** Expression of predicted miR-5352 target gene *Klf4* in ovine abomasal organoids with no stimulation, treated with IL-13 alone, control mimic + IL13 (alone, with Lipofectamine, or DharmaFECT reagent), or miR-5352 mimic + IL-13 (alone, with Lipofectamine, or DharmaFECT reagent). miRNA mimics at 100 nM final conc. **(C)** Expression of *Klf4* in ovine abomasal organoids with no stimulation, treated with IL-13 alone, control mimic + IL-13 or miR-5352 mimic + IL-13, using DharmaFECT as transfection reagent and miRNA mimic at 25, 50, 100 nM. Log2 fold change of RT-qPCR values were compared with non-stimulated control in three independent biological replicates +/−SD. Statistical analysis was by ordinary one-way ANOVA with Tukey’s multiple comparisons test; **, P < 0.01. **(D)** Representative images of abomasal control (unstimulated) organoids, stimulated with IL-13, stimulated with IL-13 + control mimic (50nM) or stimulated with IL-13 + miR-5352 mimic (50nM) at days 2, 4, 6, 8 and 12 of culture. D6 + wash shows the same treated organoids growing normally in OGM 6 days after removal of IL-13 and mimics. Scale bars: 100 µm. **(E)** Average percentage of budding organoids at day 2 after miR mimic transfection. Graph represents the average of at least 5 images and in total represents 500 organoids per treatment. Statistics calculated with a two-tailed t-test on mean of biological replicates (n=3) compared to control. **, P < 0.01. **(F)** Representative images of abomasal organoids transfected with control mimic (50nM) or with miR-5352 mimic (50nM), stimulated with IL-13 and cultured for 8 days in organoid growth media (OGM, with Rho kinase, TGF-β receptor type I and p38 MAP kinase inhibitors) or organoid differentiation media (ODM), and the same treated organoids after wash and passage in OGM alone. White arrow in ODM indicates re-grow of abomasal organoids. Scale bars: 100 µm.

Ovine abomasal organoids were then transfected with unlabelled miR-5352 mimic and gene expression examined by RT-qPCR. We focussed initially on a bioinformatically-predicted target mRNA of miR-5352 (see below), encoding Kruppel-like transcription factor KLF4. KLF4 is enriched in the GI tract and is essential for goblet cell differentiation and maintaining GI homeostasis by preventing over-proliferation of epithelial cells^44,45,46^. In the presence of IL-13, expression of *Klf4* was increased in ovine abomasal organoids relative to untreated organoids (**Figure 2B, C**). In contrast, IL-13 plus miR-5352 mimic resulted in a significant reduction in *Klf4* expression, with greatest suppression observed using DharmaFECT reagent, consistent with increased miRNA delivery (**Figure 2B, C**). Treatment of organoids with IL-13 plus control miRNA mimic (universal miRIDIAN Mimic Negative Control or a scrambled miR-5352 mimic), resulted in no suppression of *Klf4* relative to IL-13 alone (**Figure 2B, C and Supplementary Figure S2**). *Klf4* suppression occurred using a range of miR-5352 concentrations (25-100nM) and was dose–dependent (**Figure 2C**). Based on these data, all further transfections were performed using DharmaFECT reagent and parasite miRNA mimic at 50nM, a concentration similar to that used in previous studies with other miRNA mimics^47, 48^.

To test any effect of miR-5352 mimic on organoid differentiation, ovine abomasal organoids were transfected with miR-5352 or control mimic and organoid morphology examined over the course of 12 days with mimic replaced every 2-3 days. As described above, IL-13 was added to organoids to promote cell differentiation, similar to IL-13 responses to nematode infection in vivo^22^. IL-13 alone or IL-13 plus control mimic resulted in organoid budding, first observed after 48 h and increased throughout the 12-day period (**Figure 2D, E**). In contrast, in the presence of IL-13 and miR-5352 mimic, organoids failed to bud after 48 h and were similar to unstimulated organoids (control) (**Figure 2D, E**). Some budding occurred after 4 days in culture, but this was limited, and organoids started to collapse or remained smaller in size and less differentiated than those in IL-13 alone or IL-13 plus control mimic (**Figure 2D**). MiR-5352-treated organoids were still viable after 12 days; when treated organoids were washed and cultured in standard organoid growth medium (OGM, containing inhibitors to block cell differentiation) without IL-13 for six days, organoids recovered their round shape and grew normally (**Figure 2D**). These results indicate that miR-5352 can suppress differentiation of ovine abomasal organoids induced by IL-13 and, while miR-5352-treated organoids appeared abnormal, stem cells were still viable allowing organoid regrowth.

Maintenance of GI organoid stem cells following miR-5352 mimic transfection was examined further using organoid differentiation medium (ODM). This lacks inhibitors of Rho kinase, TGF-β receptor type I and p38 MAP kinase, and promotes greater cell differentiation, compared to organoid growth medium (OGM). When grown in ODM and IL-13, abomasal organoids initially showed budding but then took on a 2-dimensional appearance and stopped budding, suggesting a rapid increase in cell differentiation (**Figure 2F**). This occurred with ODM and IL-13 plus control mimic or IL-13 plus miR-5352 mimic, but not when grown in OGM and IL-13. Notably, after removal of ODM and passage in OGM, only those organoids grown in the presence of IL-13 and miR-5352 mimic were able to re-grow and differentiate/bud, while those grown in ODM and IL-13 plus control mimic did not re-grow (**Figure 2F**). This indicates that miR-5352 mimic helps maintain cell stemness, enabling organoid/tissue renewal.

### 3. miR-5352 mimic suppresses GI organoid differentiation

To examine changes in specific cell markers following abomasal organoid transfection with miR-5352 mimic, RT-qPCR was carried out. Expression of *Pou2f3* and *Klf4* was reduced, while that of stem cell markers *Olfm4* and *Lgr5* was increased relative to control mimic, in the presence of IL-13 (**Figure 3A**). These changes in expression were supported by IFA, while Ki67^+^ cell numbers were similar in the presence of control or miR-5352 mimic (**Figure 3B**). Similarly, murine SI *Dclk1* tdTomato reporter organoids stimulated with IL-13 and transfected with miR-5352 showed a reduced number of tdTomato^+^ cells relative to IL-13 plus control mimic, indicating suppression of murine tuft cell differentiation (**Figure 3C and Supplementary Figure 2**). There was also a reduction of *Klf4* and *Muc2* expression (**Figure 3D**), and a corresponding decrease in the number of KLF4^+^ and MUC2^+^ cells by IFA. Ki67^+^ cell numbers were slightly increased in the presence miR-5352 mimic in SI organoids (**Figure 3E**). These data indicate that, remarkably miR-5352 mimic alone can reduce the stimulatory differentiation effects of IL-13 on both ovine and murine GI organoids, limiting the expansion of tuft and mucous-secreting cells, while promoting stem cell maintenance.

**Figure 3.**
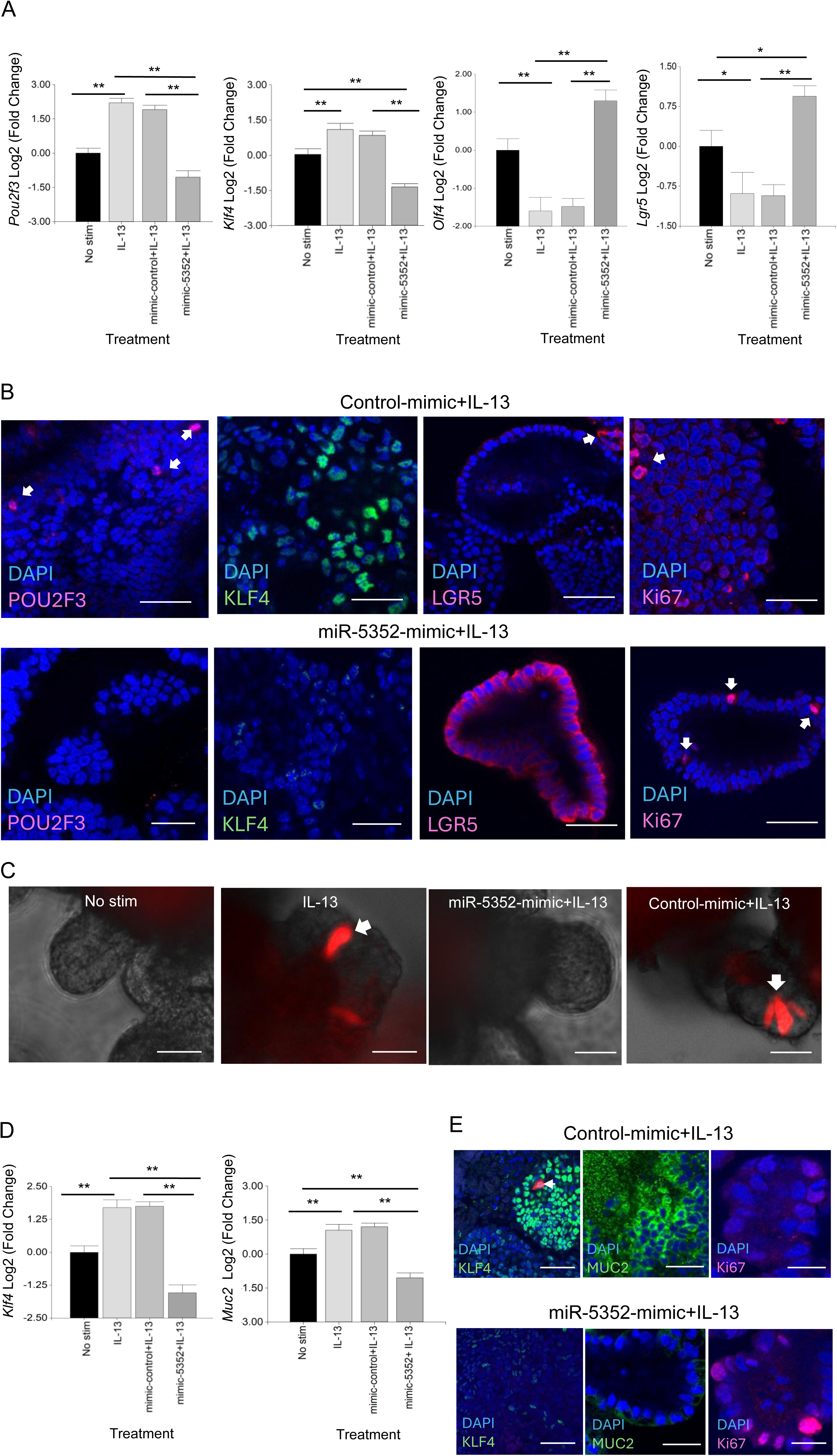
miR-5352 mimic suppresses IL-13-induced secretory cell expansion and promotes stemness. **(A)** Expression of *Pou2f3*, *Klf4, Olfm4 and Lgr5* in ovine abomasal organoids following no stimulation, IL-13 stimulation, IL-13 plus control mimic or IL-13 plus miR-5352 mimic (day 2 of culture in OGM). **(B)** Confocal microscopy showing POU2F3^+^, KLF4^+^, LGR5^+^ and Ki67^+^ cells in ovine abomasal organoids incubated with IL-13 plus control mimic or miR-5352 mimic (day 4 of culture). **(C)** Representative images of *Dclk1* tdTomato^+^ cells (arrows) in murine SI reporter organoids with no stimulation, after treatment with IL-13, IL-13 plus miR-5352 mimic or IL-13 plus control mimic (day 4 of culture). **(D)** Expression of *Klf4* and *Muc2* in murine SI organoids following no stimulation, IL-13 stimulation, IL-13 plus control mimic or IL-13 plus miR-5352 mimic (day 2 of culture). **(E)** Confocal microscopy showing KLF4^+^, MUC2^+^ and Ki67^+^ cells in murine SI organoids incubated with IL-13 plus control mimic or miR-5352 mimic (day 4 of culture). Log2 fold change of qRT-PCR values are compared with non-stimulated control in three independent biological replicates. Scale bars: 20µm. *, P < 0.05; **, P < 0.01.

### 4. Opposing effects of IL-13 and miR-5352 on global gene expression in abomasal organoids

The impact of miR-5352 on gene expression was examined at a global level. Ovine abomasal organoids alone (control), stimulated with IL-13, IL-13 plus control mimic or IL-13 plus miR-5352 mimic were subject to bulk RNA-seq and gene expression profiles compared initially by principal component analysis (PCA) (four biological replicates per condition cultured for 48 h; RNA-seq data are available at PRJNA1102187 and output data summarised in **Supplementary Table S1)**. Samples from the same treatment condition clustered together, with IL-13 inducing diametrically opposite effects on PC1 and PC2 compared with unstimulated controls with the majority of variance (98.41%) (**Figure 4A**). Gene expression profiles of samples treated with IL-13 plus control mimic overlapped with those of IL-13 alone, with a few genes showing a small change in expression (**Figure 4A** and **Supplementary Figure S3**). Notably, IL-13 plus miR-5352 mimic treated samples showed a shift in PC2. The RNA-seq data confirmed that ovine gastric epithelial cells are highly responsive to IL-13, and that the changes in gene expression induced by IL-13 are modulated by miR-5352.

**Figure 4.**
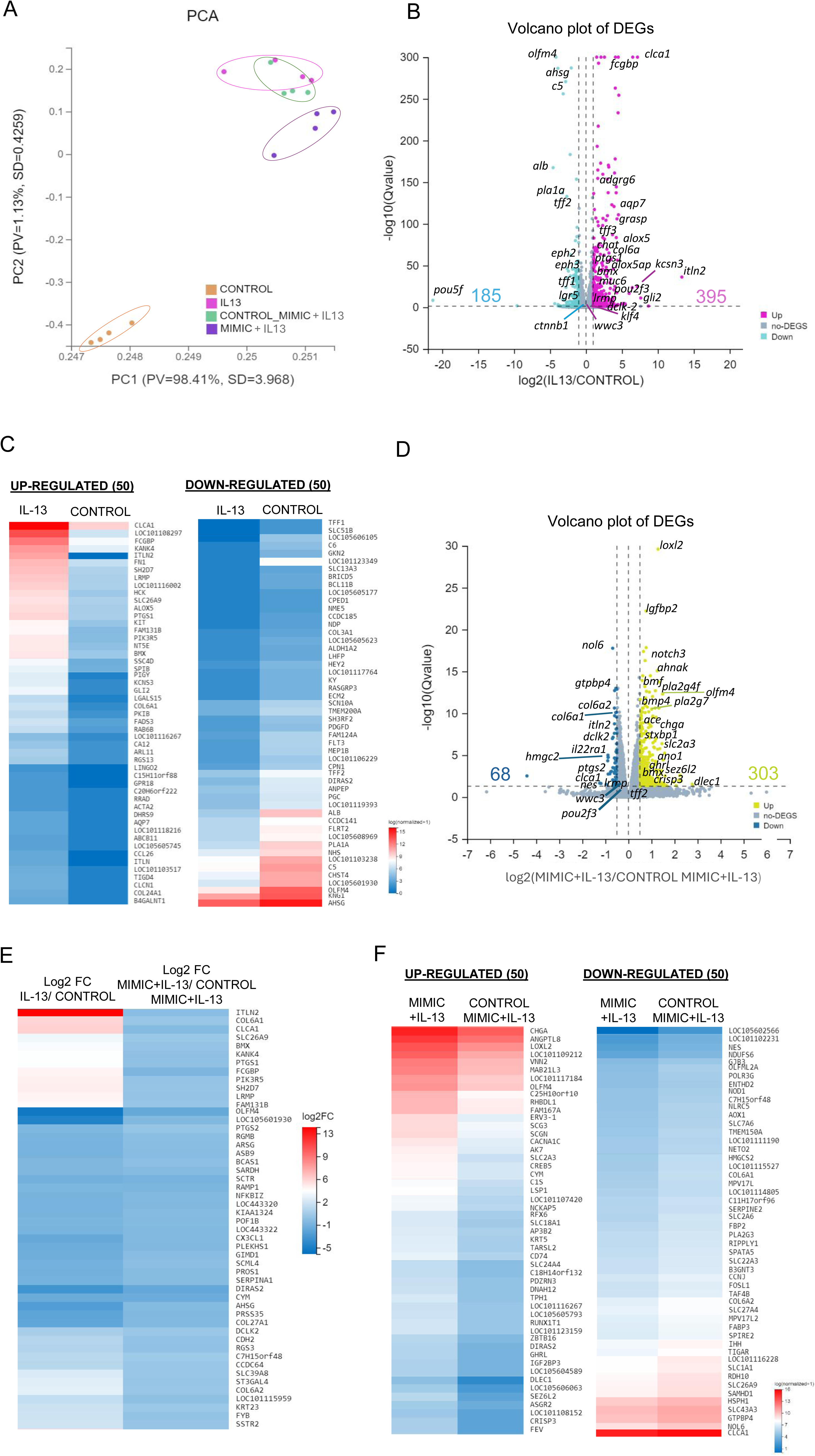
IL-13 and miR-5352 mimic have opposing effects on gene expression in ovine abomasal organoids. **(A)** Principal component analysis (PCA) plot of RNA-seq in 2 dimensions, with each dot representing a biological sample (n=4 per treatment group). PV is proportion of variance and SD is standard deviation. **(B)** Volcano plot of differentially expressed genes (DEGs), IL-13 vs. control group of mean normalized expression counts. Pink represents up-regulated DEGs, light blue represents down-regulated genes and grey represents non-DEG. **(C)** Heat map showing expression level of 50 most up- and down-regulated variant genes from IL-13 treated vs control organoids using normalised average read counts, standardized method: log(value+1). **(D)** Volcano plot of DEGs in organoids treated with IL-13 plus miR-5352 mimic versus IL-13 plus control mimic of mean normalized expression counts. Yellow represents up-regulated genes, blue represents down-regulated genes, and grey represents non-DEG. **(E)** Heat map showing expression difference (Log2 fold change) of 50 variant genes with opposite direction of regulation in IL-13/control vs miR-5352 mimic plus IL-13/ control mimic plus IL-13. **(F)** Heat map showing expression level of 50 most up and down-regulated variant genes from IL-13 plus miR-5352 mimic vs IL-13 plus control mimic using normalised average read counts, standardized method: log(value+1). In Volcano plots, x-axis represents the fold change of the difference after conversion to log2, and the y-axis represents the significance value after conversion to-log10. In Heatmaps: colours indicate level of expression from low (blue) to high (red).

We identified a total of 56,081 transcripts and 16,179 differentially expressed genes (DEGs) using a likelihood ratio test across all conditions. Treatment with IL-13 alone resulted in altered expression of 580 genes relative to unstimulated control organoids (395 up-regulated, 185 down-regulated, q-value < 0.05 and log2FC > 1) (**Figure 4B; Supplementary Table S2**). Up-regulated genes included intelectin *Itln2*, associated with type 2 responses to nematode infection^49^, *Clca1* and *Fcgbp*, expressed in mucous-producing cells, *Ptgs1*, *Alox5, Pou2f3,* and *Adgrg6*, highly enriched in ovine abomasal tuft cells based on our previous single cell (sc) RNA-seq analysis^35^, and potassium voltage-gated channel subfamily S member 3 (*Kcns3)* involved in cell cycle regulation and proliferation^50^ (**Figure 4B**). Among the most down-regulated genes in the presence of IL-13 were GI stem cell marker *Olfm4*^51^, tumour-associated glycoprotein *Ahsg*, embryonic stem cell transcription factor *Pou5f,* and intestinal stem cell niche regulators ephrin-B2 (*EphB2*) and ephrin-B3 (*EphB3)*^52^. These findings align with the role of IL-13 in promoting GI epithelial cell differentiation **(Figure 4B**). Expression profiles of the top 50 most up- and down-regulated genes are shown in **Figure 4C**.

We then evaluated the gene modulatory effects of miR-5352 mimic on IL-13-treated organoids. Comparison of transcriptional profiles of IL-13-treated ovine abomasal organoid with miR-5352 versus control mimic identified 371 DEGs (303 up-regulated and 68 down-regulated; q-value < 0.05 and log2FC > 0.5) (**Figure 4D** and **Supplementary Table S3**). The high number of DEGs most likely indicates secondary effects of miR-5352 on the organoids, in addition to direct regulation of miRNA target genes. Up-regulated genes included *Loxl2* involved in senescence and tumour suppression^53^, insulin-like growth factor *Lgfbp2*, *Notch3*, involved in maintaining stemness of cancer cells^54^ and *Olfm4*. Down-regulated genes included *Nol6* nucleolar protein that stimulates cell growth and gastric cancer^55^, GTP binding protein 4 *gtpbp4,* involved in cell cycle regulation, *Itln2,* tuft cell genes *Dclk2 and Ptgs2* and mucous cell gene *Clca1.* Notably, 390 genes (q-value < 0.05) showed the opposite direction of regulation (opposite Log2FC) in abomasal organoids treated with IL-13 versus no treatment, compared to IL-13 plus miR-5352 mimic versus IL-13 plus control mimic (**Figure 4E** and **Supplementary Table S4**). These included *Itln2*, *Clca1, Olfm4*, and tuft cell genes *Bmx, Ptgs1* and *2, Lrmp* and *Dclk2*, indicating that the parasite miRNA is reversing some of the IL-13 stimulatory effects on organoid gene expression, especially on tuft, mucous and stem cells (**Figure 4E**). Expression profiles of the top 50 most up- and down-regulated genes in abomasal organoids treated with IL-13 plus miR-5352 relative to IL-13 plus control mimic are shown in **Figure 4F**.

### 5. miR-5352 promotes stemness and represses tuft and mucous cell gene set expression in ovine abomasal organoids

To describe the functional features of abomasal organoids treated with IL-13 or IL-13 plus miRNA mimic we implemented a gene set enrichment analysis (GSEA) on DEGs for Biological Process domain (GO:BP) terms^56^. Consistent with the observed stimulatory effect on organoid cell differentiation, treatment with IL-13 caused a marked increase in normalized enrichment score (NES) associated with G1/S transition of mitotic cell cycle, stem cell differentiation, negative regulation of Notch signalling pathway which promotes renewal of intestinal and gastric stem cells^57,58^ (**Figure 5A**), regulation of cell shape and positive regulation of cell differentiation, among others **(Supplementary Table S5**). Conversely, GSEA for abomasal organoids treated with IL-13 plus miR-5352 mimic showed enrichment of terms associated with negative regulation of G1/S transition of mitotic cell cycle, cell cycle arrest, positive regulation of Notch signalling pathway (**Figure 5B**) and negative regulation of cell growth (**Supplementary Table S6**).

**Figure 5.**
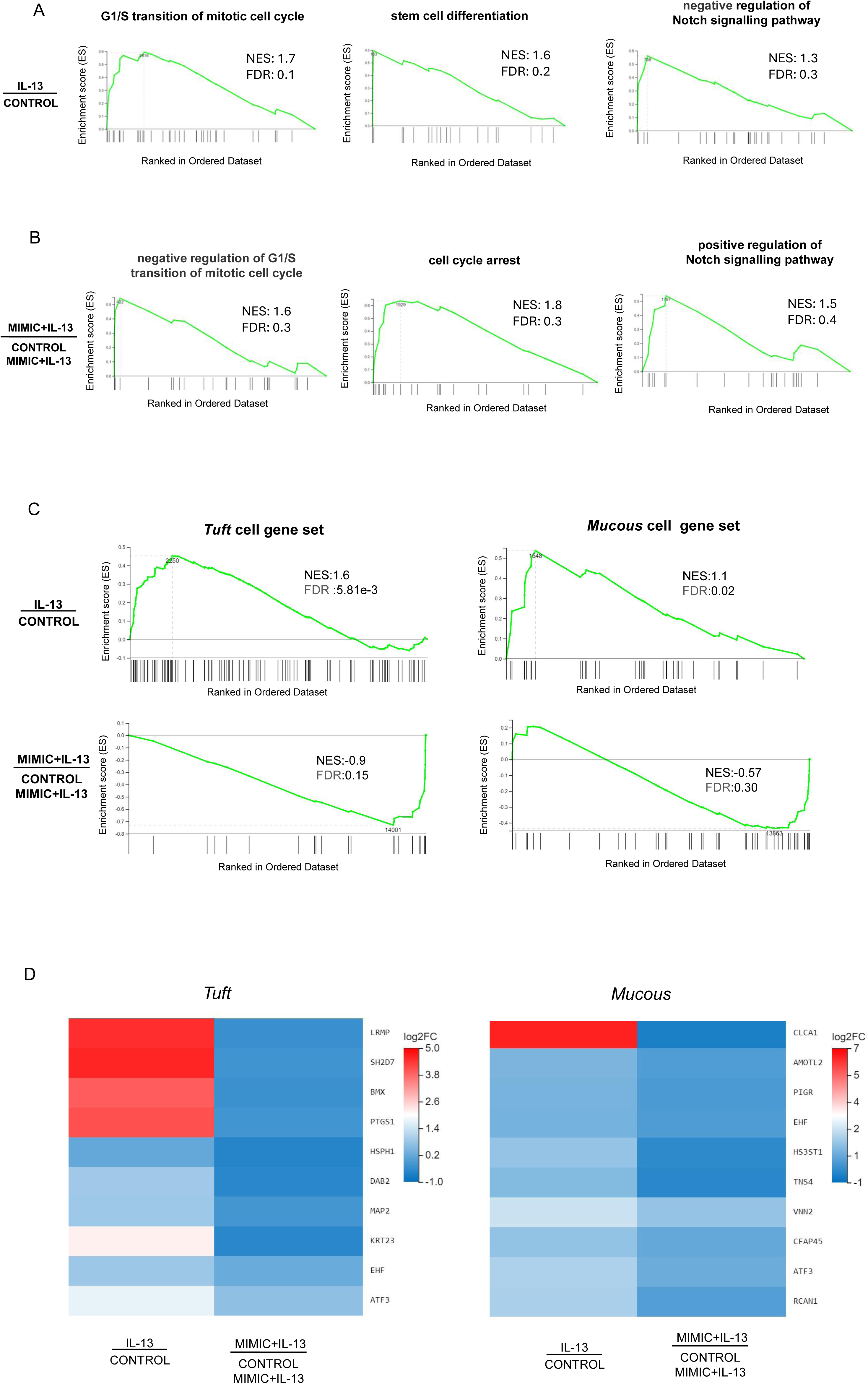
Gene sets associated with differentiation are enriched by IL-13 and suppressed by miR-5352. **(A)** Gene set enrichment analysis (GSEA) of genes expressed in ovine abomasal organoids treated with IL-13 vs. no treatment (control). Graphs depict the enrichment score (ES) on the y-axis, with positive values where gene sets are induced, and negative values where they are inhibited. Normalised enrichment score (NES) and false discovery rate (FDR) are indicated on each graph. Each black vertical line represents one gene in category GO:BP and its relative ranking against all genes analysed. **(B)** GSEA of genes expressed in organoids treated with IL-13 plus miR-5352 mimic vs. IL-13 plus control mimic. **(C)** GSEA of tuft or mucous cell gene sets (Hildersley et al., 2021) expressed in abomasal organoids exposed to IL-13 vs no treatment (control) and miR-5352 mimic plus IL-13 vs control mimic plus IL13. Each vertical bar on the x-axis represents an individual gene within the gene set for the stated cell type. **(D)** Heat map of tuft and mucous cell genes showing opposite direction of regulation (log2 fold change) in organoids treated with IL-13 vs. no treatment (control) and miR-5352 mimic + IL-13 vs. control mimic + IL13.

Additionally, using tuft and mucous gene sets identified by our previous scRNA-seq of abomasal mucosa^35^, GSEA identified an enrichment for these genes when organoids were stimulated with IL-13 vs no treatment, and a reduction when organoids were treated with IL-13 plus miR-5352 compared to IL-13 plus control mimic (**Figure 5C**). Multiple tuft cell canonical genes, such as *Lrmp, Bmx, Ptgs1*, and mucous cell genes *Clca1, Amotl2, Tns4*, were suppressed by miR-5352 mimic treatment, even in the presence of IL-13 (**Figure 5D**). Reduced NES scores for these cellular gene sets further supports suppression of GI secretory cell differentiation by miR-5352.

### 6. *H. contortus* secreted miRNAs are predicted to target host signalling pathways and IL-22 response

Regulatory targets of miRNAs can be predicted in silico using bioinformatic programmes^59,60^. We applied three programmes (MiRanda, RNAhybrid and TargetScan) to predict target mRNAs of Hco-miR-5352 and the other 22 miRNAs identified in *H. contortus* ES^34^. Merging bovine and ovine bioinformatic target prediction datasets for Hco-miR-5352 identified 54 potential target mRNAs, with 26 of these identified by at least two programmes, giving more confidence to the prediction data (**Figure 6A**, **Supplementary Table S7**). Fifty-one of the Hco-miR-5352 predicted target genes were also found to be targeted by other miRNAs present in *H. contortus* ES, including Hco-miR-5960, Hco-lin-4, Hco-miR-61, Hco-miR-43 and Hco-miR-5895. The latter three miRNAs are members of the Hco-miR-5352 cluster conserved across GI nematodes^34^, suggesting that additional conserved parasite miRNAs are potentially involved in co-regulating host genes and may produce additive effects. Seven predicted target mRNAs, down-regulated in the presence of miR-5352 mimic from our abomasal organoid RNA-seq data, were predicted as hco-miR-5352 targets in only one of the two host species genomes; miRNAs binding site were manually identified in the 3′UTR, including for *Il22ra1* (**Supplementary Table S7**).

**Figure 6.**
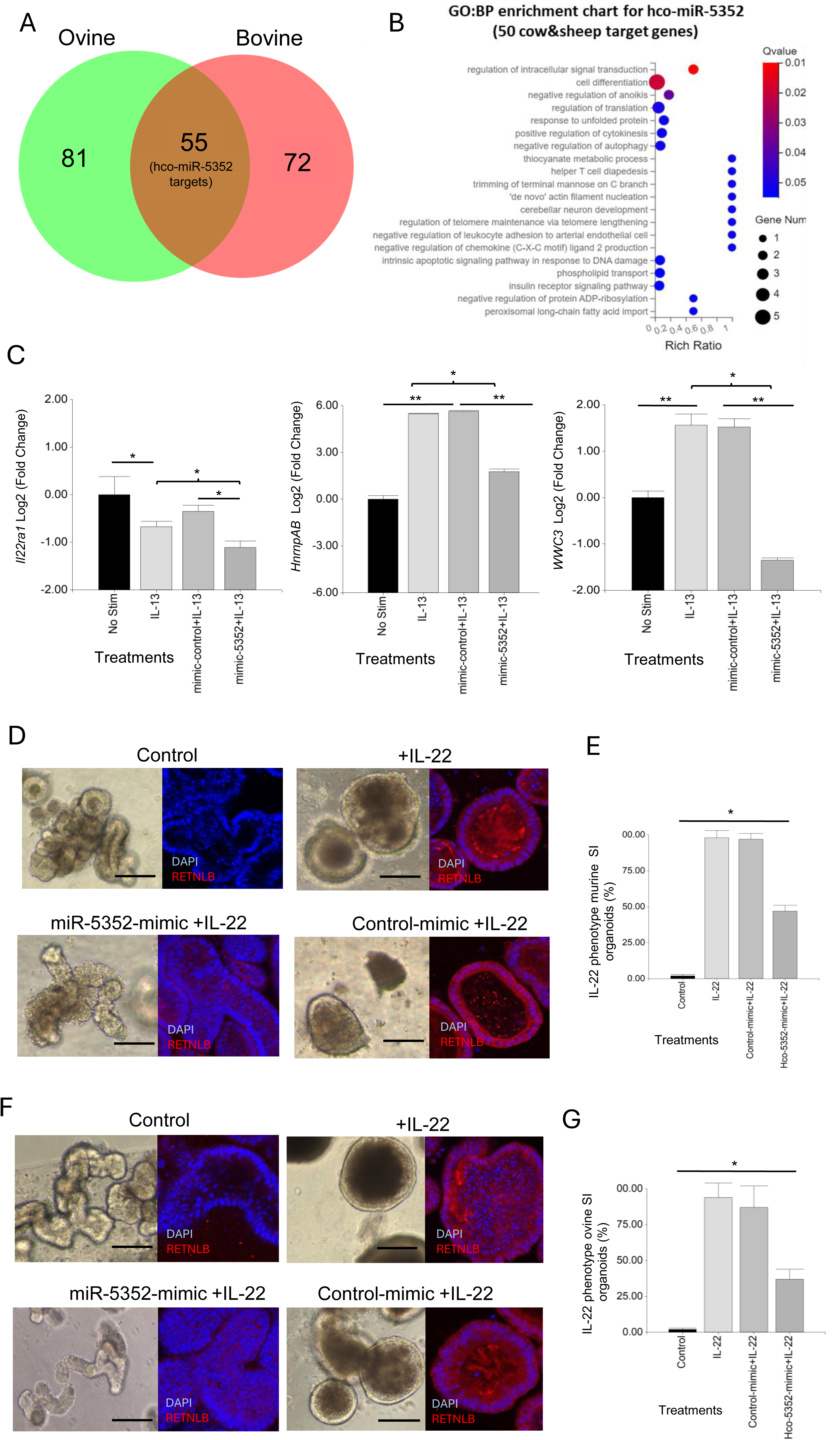
Hco-miR-5352 predicted target genes include host signalling pathways. **(A)** Venn diagram of hco-miR-5352 predicted targets from ovine and bovine bioinformatic genome predictions. **(B)** GO: Biological Process enrichment bubble chart of the 49 predicted target genes of Hco-miR-5352 in bovine and ovine genomes and expressed in ovine abomasal organoids by RNA-seq (q-value < 0.05). **(C)** RT-qPCR of selected hco-miR-5352 predicted target mRNAs from GI organoids following no treatment or treated with IL-13, IL-13 plus control mimic or IL-13 plus hco-miR-5352 mimic. *Il22ra1* was amplified from murine SI organoids, *WWC3* and *hnrnpAB* were amplified from ovine abomasal organoids, all at day 2 after treatment. Log2 fold change of RT-qPCR values are compared with non-stimulated control in three independent biological replicates. *, P < 0.05; **, P < 0.01. **(D)** Representative confocal microscopy images of murine SI control (untreated) organoids, treated with IL-22, or IL-22 plus control mimic (50nM) or IL-22 plus miR-5352 mimic (50nM) at day 4 after treatment, showing RELMB (Resistin-like beta, RETNLB) IFA. **(E)** Average percentage of IL-22 phenotype (small, non-budding, dark organoids) in murine SI organoids at day 4 after no treatment, or treatment with IL-22, IL-22 plus control mimic or IL-22 plus miR-5352 mimic. **(F)** Representative confocal images of ovine SI control (untreated) organoids, treated with IL-22, treated with IL-22 plus control mimic (50nM) or treated with IL-22 plus miR-5352 mimic (50nM) at day 4 after treatment **(G)** Average percentage of IL-22 phenotype in ovine SI organoids at day 4 after control, IL-22, IL-22 plus control mimic or IL-22 plus miR-5352 mimic treatment. Graphs represent the average of at least 5 images and in total represents 500 organoids per treatment. Statistics calculated with a two-tailed t-test on mean of biological replicates (n=3) compared to control. Scale bars: 300 µm.

Of the 61 total predicted Hco-miR-5352 targets, six were not found to be expressed in abomasal organoids by our bulk RNA-seq (Cluster of Differentiation 69 (*CD69*), nuclear receptor binding protein 2 (*Nrbp2*), Orthodenticle Homeobox 2 (*Otx2*), Alanyl-tRNA synthetase 1 (*Aars1*), cytochrome P450 family 4 subfamily V polypeptide 2 (*Cyp4v2*) and Coenzyme Q8A (*Coq8a*)) (**Supplementary Table S7**). The remaining predicted targets showed enrichment (q-value <0.05) for GO:BP terms regulation of signal transduction and cell differentiation (**Figure 6B**). From our ovine abomasal RNA-seq data, 23 of the 55 predicted targets (42%) were down-regulated following transfection with miR-5352 mimic plus IL-13 relative to control mimic plus IL-13. These down-regulated targets included: *Il22ra1*, encoding the receptor for cytokine IL-22 which modulates expression of mucins in the intestinal mucous layer^61^ and promotes Paneth cell maturation by suppressing Wnt and Notch signalling pathways^62,63^; *hnrnpAB* (heterogeneous nuclear ribonucleoprotein A/B), which inhibits Notch signalling^64^; and *WWC3,* which inhibits Wnt and promotes Hippo signalling^65^ **(Supplementary Table 7)**. RT-qPCR supported the RNA-seq data, with all three transcripts reduced in GI organoids 2 days after treatment with miR-5352 mimic plus IL-13 compared to IL-13 treatment alone or IL-13 plus control mimic (**Figure 6C**). *Il22ra1* proved difficult to detect in ovine abomasal organoids. However, this transcript could be amplified from murine SI organoids, suggesting higher *Il22ra1* expression in SI than in gastric epithelia. In contrast to *hnrnpAB* and *WWC3*, which increased following IL-13 treatment, expression of *Il22ra1* decreased with IL-13 and decreased further following treatment with IL-13 plus miR-5352 mimic vs IL-13 plus control mimic (**Figure 6C**).

To test any reduction in response to IL-22 in the presence of miR-5352 mimic, murine and ovine SI organoids, were treated with IL-22 and organoid morphology was examined over the course of 4 days. IL-22 alone or IL-22 plus control mimic resulted in a change in organoid phenotype relative to untreated control organoids, first observed after 12 h. This phenotype was characterised by reduced budding, a darker appearance and an increase of the goblet cell marker RELM-β (Resistin-like beta molecule) (**Figure 6D, F**), strongly induced by IL-22 ^66,67^. In contrast, in the presence of IL-22 and miR-5352 mimic, 53% of the murine SI organoids (**Figure 6D, E**) and 63% of ovine SI organoids (**Figure 6F, G**) failed to show this IL-22 phenotype after 4 days and were similar to untreated organoids (control). These results, together with target prediction and RT-qPCR data, suggest that miR-5352 can suppress IL-22 effects by reducing expression of IL-22RA1 on epithelial cells.

### 7. miR-5352 suppresses *Klf-4* and mimics a host regulatory mechanism

Another high confidence predicted target of hco-miR-5352 is the previously mentioned Kruppel-like transcription factor gene *Klf4,* which was down-regulated following miR-5352 mimic transfection of abomasal organoids, by RT-qPCR (**Figures 2B and C);** the read count for *Klf-4* was also decreased by miR-5352 in our RNA-seq data, but this was not significant (**Supplementary Table S7**). KLF4 can bind to the transcriptional activation domain of β-catenin, inhibiting β-catenin-mediated transcription of Wnt-responsive genes, thus promoting cell differentiation^68^ (**Figure 7A**). The expression of β-catenin (*CTNNB1*) was examined by RT-qPCR in abomasal organoids at 2 and 4 days after treatment with IL-13 alone, or IL-13 plus miR-5352 or control mimic. Levels were increased with miR-5352 mimic plus IL-13 relative to controls and were similar to unstimulated abomasal organoids (**Figure 7B**). These changes in expression were supported by CTNNB1 IFA in ovine abomasal (**Figure 7C**) and ovine SI (**Figure 7D**) organoids. In some cells, nuclear localisation of CTNNB1 was evident, suggesting cytoplasmic-nuclear shuttling of β-catenin and activation of the Wnt/β-catenin pathway in the presence of miR-5352 mimic (**Figure 7C, D**). This implies that miR-5352 mitigates the IL-13-mediated down-regulation of *CTNNB1* (**Figure 7B and Supplementary Table S2**), potentially through direct suppression of *Klf4* expression. Notably, recovery of *CTNNB1* level was detected at day 4 after treatment, but not at day 2, suggesting a lag time between a reduction in *Klf4* and increase in *CTNNB1.* Our target prediction data indicate that miR-5352 potentially modulates expression of multiple target genes involved in Wnt and Notch regulation: *Klf4, Il22ra1, hnrnpAB* and *WWC3*. From RNA-seq data, the miRNA-mediated down-regulation of these predicted targets is modest, as may be expected for miRNA activity^69^. Additively, the overall effect is inhibition of tuft and mucous cell differentiation and promotion of stemness, via modulation of Wnt and Notch signalling.

**Figure 7.**
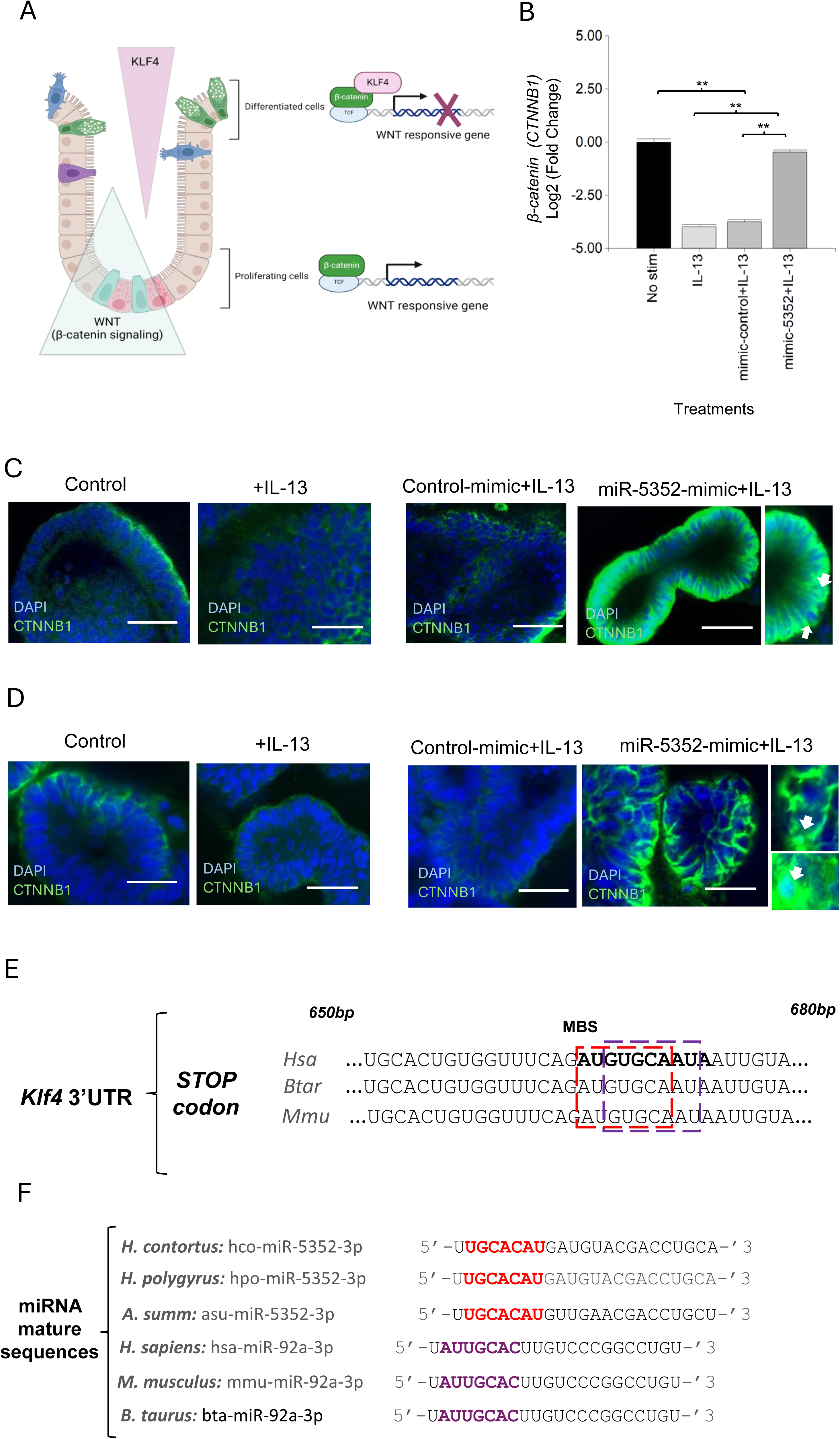
miR-5352 mediates suppression of *Klf-4* Wnt signaling and can mimic a host regulatory mechanism. **(A)** Model for KLF4-mediated regulation of β-catenin (based on Ref 68.) KLF4 expressed in differentiated epithelial cells binds to β-catenin and suppresses β-catenin-dependent Wnt signalling, resulting in differentiation. Down-regulation of KLF4 relieves β-catenin suppression and promotes Wnt signalling and cell proliferation. **(B)** Expression of β-catenin (*CTNNB1*) in ovine abomasal organoids treated with IL-13, IL-13 plus control mimic or IL-13 plus miR-5352 mimic (day 4 after treatment). Log2 fold change of qRT-PCR values are compared with non-stimulated control in three independent biological replicates. *, P < 0.05; **, P < 0.01. **(C)** Confocal IFA microscopy showing CTNNB1^+^ cells in abomasal organoids untreated or incubated with IL-13 alone, IL-13 plus control mimic or miR-5352 mimic (day 4 of culture). Scale bars: 20µm. Arrows indicate CTNNB antibody localisation in the nuclei of epithelial cells. **D)** Confocal microscopy showing CTNNB1^+^ cells in ovine SI organoids untreated or incubated with IL-13 alone or IL-13 plus control mimic or miR-5352 mimic (day 4 of culture). Scale bars: 20µm. **(E)** Overlap of predicted miRNA binding site (MBS) for Hco-miR-5352 (red) and Hsa-miR-92a (purple) in *Klf4* 3’UTR from human, bovine and murine. MBS is approximately 670bp downstream of the stop codon (miRanda and RNAhybrid data). **(F)** Alignment of seed sequence (nucleotides 2-8) of miR-5352 sequences from GI nematode species (red) with mammalian (human, murine, bovine) miR-92a (purple). MirBase accession numbers: hco-miR-5352-3p, MI0020053; hpo-miR-5352-3p, MI0032733; asu-miR-5352-3p, MI0018601 (*Ascaris suum*); hsa-miR-92a-3p, MIMAT0000092; mmu-miR-92a-3p, MIMAT0000539; bta-miR-5352-3p, MIMAT0009383. MiR-5352 sequences identified in additional GI nematodes are detailed in Ref 34.

In humans, expression of *Klf4* is regulated post-transcriptionally by Hsa-miR-92a^70,71^. By analysing the *Klf-4* 3’UTR sequence, we found that the predicted binding site for miR-5352 overlaps with that for Hsa-miR-92a and is conserved across humans, mice and ruminants (**Figure 7E**). The seed sequence (nucleotides 2-8) of Hco-miR-5352 partially aligns with mammalian miR-92a seed and is conserved across different GI nematodes (**Figure 7F**). This suggests that miR-5352 in GI nematodes may have evolved to mimic host miR-92a and can phenocopy effects on GI tissues, promoting cell stemness and reducing differentiation^71^.

## Discussion

Parasitic helminths are remarkable in establishing persistent infections within their hosts, often eliciting minimal inflammatory responses and employing a range of mechanisms to prevent or delay their expulsion^72^. Here we report that ES products from *H. contortus* can suppress effects of the type-2 cytokine IL-13 on tuft and mucous cell differentiation in ovine and murine GI organoids. Remarkably, a single miRNA, miR-5352, present in ES from *H. contortus, H. polygyrus* and other GI nematodes^12,34^ achieved the same effect. We identified target mRNAs down-regulated following miRNA mimic transfection of organoids, and our data indicate that parasite-mediated enhancement of Wnt and Notch signalling are key mechanisms of miR-5352-mediated immunomodulation in host GI cells.

Helminth miRNAs have been sequenced from secreted EV or non-vesicular ES products^12,15,16,17,18^ and their interaction with host cells identify them as potential regulators of host gene expression. EVs released by *H. polygyrus* adult worms can down-regulate expression of host *Il33r* and *Dusp1* in macrophage cells in vitro^12^. Several EV miRNAs were proposed to be involved, but the precise mechanisms are not yet elucidated. The focus of this study, miR-5352, is enriched in non-vesicular ES released from *H. contortus* parasitic L4 larvae and adult stages, with lower abundance in EV^34^. Notably, we show that the seed sequence of GI nematode miR-5352 is similar to host miR-92a (miR-25 family) and that the miRNAs have overlapping binding sites in the 3’UTR of target mRNA *Klf4.* Human miR-92a regulates host cell proliferation^73^ and maintains stemness in the GI tract^74^, consistent with our observed effects of miR-5352 mimic on GI organoids. In support of our findings, previous studies reported that miRNAs in helminth ES show sequence similarity to host miRNA sequences, including miR-100, let-7, lin-4 and bantam^12,14,34^. Helminth secreted miRNAs may, therefore, represent a form of molecular mimicry to aid parasite survival, analogous to parasite proteins such as *H. polygyrus* TGF-β mimics which can mimic the biological functions of host TGF-β^75^. From an evolutionary perspective, the conservation of miRNA and 3’UTR regulatory sequences across host and parasite suggests that mutational changes and potential host ‘resistance’ to parasite manipulation may be unlikely to occur^76^, and that cross-species regulation may have arisen as parasites and hosts co-evolved towards a tolerance relationship.

The modulatory effects of hco-miR-5352 may be enhanced by the action of other *H. contortus* secreted miRNAs, including those from the same cluster, and/or other ES components, such as other small RNAs or proteins. Indeed, we found that some of the predicted targets of hco-miR-5352 could potentially be targeted by other miRNAs in ES of *H. contortus* and other helminths. As well as suggesting additive regulatory effects, this lends confidence to the predictions, since 3’UTRs with more than one target site are more likely to be genuine miRNA targets^77^.

Additionally, our previous studies detected hco-miR-5352 (and other hco-miRs) in abomasal mucosa and mesenteric lymph nodes following *H. contortus* infection^34^, suggesting that *H. contortus* miRNAs may modulate expression of epithelial and immune-associated genes. Further studies employing immune cells alone or co-culture of organoids with immune cells will help determine additional regulatory effects of helminth ES and miRNAs on host immune function and epithelial-immune cell cross-talk.

A number of miR-5352 targets identified here (*Klf4, WWC3, hnrnpAB, Il22ra1*) are involved in regulating Wnt and/or Notch signalling. MiR-mediated down-regulation of these target genes is predicted to enhance Wnt/Notch signalling, promoting GI epithelial cell stemness and reducing secretory cell differentiation. Drurey et al.^22^ previously proposed that the increase in *Hes1* following exposure of murine SI organoids to *H. polygyus* ES suppressed *Atoh1*, and subsequently secretory cell numbers, however the mechanism of *Hes1* induction was not established. Our data suggest that miR-5352-driven suppression of target genes enhances Notch and expression of its downstream gene *Hes1*. Suppression of *Il22ra1* by miR-5352 is a potential additional mechanism by which nematodes may modulate epithelial cell differentiation. IL-22, produced mainly by Th1, Th17 and ILC3 cells^78^, promotes mucous/goblet cell hyperplasia, and production of the antimicrobial peptide RELM-β involved in the weep and sweep response^67,79^. IL-22 deficient mice show impaired expulsion of the SI nematode *N. brasiliensis* and colonic whipworm *T. muris*, despite strong type-2 cytokine responses^79^. In a more recent study^80^, IL-22 was found to synergise with IL-1β in production of antimicrobial peptides and neutrophil chemoattractant cytokine CXCL1 produced by ileum and colon epithelial cells, to more effectively clear *Citrobacter rodentium* infection. A reduction in neutrophil infiltration, resulting from reduced IL22RA1 levels, may be an additional mechanism enabling GI nematode survival.

Regulation of *Il22ra1* by miR-92a and/or other miRNAs, may have evolved as a host feedback mechanism to limit mucous cell expansion. Recent studies have identified additional novel regulatory mechanisms for control or resolution of host GI immune responses and inflammation: BMP signalling induced by IL-13 was shown to restrict the increase in tuft cell numbers following IL-13 treatment or *N. brasiliensis* infection^67^, while the IL-25 receptor IL-17RB present on tuft cells acts as a brake on their production of IL-25 that stimulates ILC2 cells^81^. Multiple mechanisms are likely at play to limit innate and adaptive responses potentially associated with immunopathology or inappropriate activation, and these mechanisms may have been exploited by nematodes for amelioration of immune defences.

Abomasal or SI organoids exposed to *H. contortus* ES or miR-5352 mimic alone showed relatively normal organoid growth morphology, albeit with less differentiation of secretory cells, and maintained a stem cell population capable of regrowth/renewal. This contrasts with the spheroid morphology observed after treatment of murine SI organoids with concentrated ES of *H. polygyrus*^22,42^ or ES of *Ostertagia ostertagi* L3 larvae^82^. Spheroid or foetal-like regions of the SI were also reported following *H. polygyrus* infection^43^. Spheroid formation is likely to be an extreme and localised effect of high concentration of ES, which interferes with both cell differentiation and renewal. While this response is not unique to *H. polygyrus*, ES from this nematode appears to have a more potent effect than *H. contortus* ES. As each helminth species has a different predilection site within the host, some differences between parasite ES effects may be expected. Detailed comparisons of ES components across species may be predicted to identify conserved and unique molecule(s) involved. Notably, data in Karo-Atar et al.^42^ show that *H. polygyrus* ES also reduced expression of *Il22ra1* and *Klf4* in SI murine organoids, and increased expression of *Lgr5* and *Olfm4*, although a causative mechanism was not proposed. Our findings suggest that this can be mediated by nematode miR-5352 alone, and may act in combination with other ES components, which can be examined in future studies.

Specific target genes and pathways regulated by helminth miRNAs can help reveal novel therapeutic approaches for GI parasite control and for immune- or damage-related disorders^83^. “Helminth therapy” can be applied to autoimmune and chronic inflammatory disorders of the GI tract responsible for inflammatory bowel disease (IBD)^84^ but the molecular mechanisms involved in reducing inflammation or repairing damaged tissue are unclear. We show that a single GI nematode secreted miRNA can regulate host genes to promote stem cell maintenance, regeneration of host GI epithelial tissue and potentially reduce inflammation and immune cell migration. Therapeutic intervention using miRNAs is an established approach that can modulate multiple gene networks using stable oligonucleotides to enhance (miRNA mimic) or interfere with (anti-miR) endogenous miRNAs^85^. Notably, higher levels of IL-22 are associated with Crohn’s disease^86^, thus miRNA-mediated suppression of *Il22ra1* could help alleviate disease symptoms. Future studies testing delivery and stability of mimics and inhibitors of miR-5352 and other nematode miRNAs, will help evaluate their potential to influence infection outcome and treat immune-related disorders.

## Materials & Methods

### Animal and parasite samples

Animal procedures were performed at Moredun Research Institute (MRI) under license as required by the UK Animals Scientific Procedures Act 1986, with ethical approval from MRI Animal Experiments Committee. Ovine abomasum (gastric) tissues for organoid cultures were obtained from four different naïve animals raised at MRI under conditions designed to exclude accidental infection with helminth parasites. Tissue was removed using a sterile scalpel and placed into sterile ice-cold Hank’s buffered saline solution (HBSS; Sigma) to be transported to the laboratory.

Adult *H. contortus* parasites, MHco3(ISE) strain, were obtained from three different male lambs infected with 5,000 *H. contortus* third stage (L3) larvae. Adults were collected at post-mortem at day 42 post-infection (p.i). The abomasum was opened and washed with 0.9% saline solution to remove and collect the worms, which were then extensively washed in saline solution and transported to the laboratory in HBSS.

### Culture of ovine abomasum and small intestine 3D organoids

Gastric glands were obtained from the fundic gastric fold of ovine abomasum. The gastric glands were extracted, and organoid cultures established from gland stem cells as described by Smith et al.^87^. Briefly, gastric glands were resuspended in 1 ml advanced DMEM/F12 (12634-010; Gibco) containing 1X B27 supplement minus vitamin A (12587-010; Gibco), 25 µg/ml gentamicin and 100 U/ml penicillin/streptomycin and counted using 10 µl of the suspension under the microscope in a Petri dish. Between 300 to 350 gastric glands per well were seeded into 24-well flat-bottomed plates (Corning) using 50 µl of Matrigel (Corning). After incubation for 20 min at 37°C, 650 µl of IntestiCult Organoid Growth Media (OGM) (STEMCELL Technologies, 6005) with 50 µg/ml gentamicin was added. To maintain organoid stemness, inhibitors were added: i) selective Rho kinase inhibitor: Y-27632 (10µM) (Cambridge Bioscience, 10005583-1 mg-CAY) ii) selective TGF-β receptor type I (TGF-βRI) kinase inhibitor:LY2157299 (500nM) (Cambridge BioScience, 15312-1 mg-CAY) and iii) selective p38 MAP kinase inhibitor: SB202190 (10 µM) (Enzo Life Science, ALX-270-268-M001) as previously reported by Smith et al.^87^. Organoids were established and grown at 37°C in 5% CO_2_ with a change of medium every 2-3 days. Organoids were passaged three times by dissociation and reseeded at 300 to 350 organoids per well in 50 µl Matrigel (Corning®, CLS356231-1EA), prior to experimental treatments. For specific experiments to increase stem cell differentiation, IntestiCult™ Organoid Differentiation Medium (ODM) (Catalog # 100-0214) was used. Ovine small intestinal (SI) organoids were provided by Dr David Smith (MRI) and cultured under the same conditions as for abomasum organoids.

### Murine small intestine tuft cell reporter 3D organoids

Organoids from mice combining conditional (Cre-dependent) tdTomato reporter allele known as Ai14 (Gt (ROSA)26Sortm14(CAG-tdTomato)Hze; JAX:007908) and inducible, tuft cell-specific, Cre enzyme (Dclk1-Cre^ERT2^) alleles were generated at University of Montpellier by PJ, FH, FG. Details of the Dclk1-Cre^ERT2^ allele (Dclk1^tm1.1(cre/ERT2)Jay^) are available from Mouse Genome Informatics (MGI) (https://www.informatics.jax.org/allele/MGI:5563428).

Tamoxifen was added at final concentration of 0.5 μM for one day to induce recombination. Organoids were cultured in IntestiCult OGM with 50 µg/ml gentamicin, at 37°C in 5% CO_2_. The medium was changed every 2-3 days, and organoids were passaged three times by dissociation and reseeding before experimental treatments^88^. The phenotype of organoids was checked using an EVOS M5000 Imaging System (Thermo Fisher Scientific) three days after tamoxifen treatment.

### Preparation of *H. contortus* excretory-secretory (ES) products

One hundred *H. contortus* adults (50 males and 50 females) were collected from three different infected animals and washed six times in warm HBSS. Further washes were carried in HBSS plus penicillin/streptomycin (2U/ml), followed by washing for 20 minutes in HBSS gentamicin (0.5 μg/ml) then worms were rewashed four more times with HBSS. Finally, worms were cultured in 50 ml of HBSS without phenol red (Sigma H864) at 37°C in 5% CO2. Medium containing ES products was collected at 24 h, centrifuged at 1500 g for 5 minutes and filtered through a 0.22 μm filter to remove released eggs and hatched L1 stages from adult ES. The ES was stored at −80°C until use.

*H. contortus* ES was used neat or was concentrated using Vivaspin 2, membrane of 3,000 MWCO PED following the manufacturer’s instructions. *H. polygyrus* ES (1.5 mg/ml) was prepared as previously described by Johnston et al.^89^. Qubit® Protein Assay Kit was used for protein quantitation of ES following the manufacturer’s instruction. Concentrated ES was used at 10 µg/ml and neat ES at 2 µg/ml final concentration. The protein concentration for all conditions is shown in **Supplementary Table S8**.

### MiRNA mimic uptake assay

To optimize the uptake of miRNA mimics into organoid cells, preliminary transfection experiments were performed using Cy3-labeled Hco-miR-5352 mimic at a final concentration of 100 nM. Transfection was performed using i) Cy3-labeled Hco-miR-5352 mimic mixed with 300 organoids and incubated for 1 hr at 37°C in 5% CO2, ii) Cy3-labeled Hco-miR-5352 mimic mixed first with lipofectamine TM 3000 reagent according to the manufacturer’s instructions, iii) Cy3-labeled Hco-miR-5352 mimic mixed with DharmaFECT TM according to the manufacturer’s instructions. Organoids were resuspended in advanced DMEM/F12 and plated in 30 μl of Matrigel into 8-well chamber-slide with 400 μl of IntestiCult OGM media for 12 h at 37°C in 5% CO2. For imaging, the organoids were prepared as described in O’Rourke et al. (2016) with some modifications. Briefly, the culture media was removed and replaced with ice-cold 10% buffered formalin from Cell Path (BAF-6000-08A). For fixation, samples were kept at room temperature for 20 minutes, washed twice with immunofluorescence (IF) buffer (0.1% Tween20 in PBS), then 300 μl of DAPI (1 μg/ml) were added for 5 min. The wells were washed three times with 300 μl IF buffer. Finally, slides were mounted using ProLong Gold antifade mountant (P10144, Thermo Fisher Scientific) and imaged by inverted confocal microscopy using a Zeiss LSM 710 Zeiss Zen Black operating software.

### *H. contortus* miRNA mimics and transfection into GI organoids

To study effects of miR-5352 on host epithelial cell gene expression, we used double-stranded RNA oligonucleotide mimic (miRIDIAN microRNA Mimic-Dharmacon^TM^) at a final concentration of 50 nM (miR-5352 mimic). The final concentration was selected by testing the dose–dependent effects of miRNA mimics (25, 50 and 100 nM) on predicted target gene expression^47^. The universal miRIDIAN Mimic Negative Control – DharmaconTM (Catalog Item: CN-0001000-01-50) and scrambled version of miR-5352 (DharmaconTM), both at final concentration of 50nM and with identical design and modifications as miRIDIAN miR-5352 mimic, were used to determine miR-5352 sequence-specific effects (sequences of miRNA mimics shown in **Supplementary Table S9**).

Organoids were treated with 400 ng/ml of IL-13 (for ovine abomasum or SI organoids, recombinant ovine IL-13, Kingfisher Biotech, INC; or for murine SI organoids, recombinant murine IL-13, PeproTech 210-13) to increase the number of secretory epithelial cells, similar to natural infection, as performed in Drurey, et al.^22^. In our experiments, IL-4 was not added since it shares the receptor subunit IL-4Rα with IL-13^19^. Organoids without IL-13 were used as controls. To study the effect of miR-5352 mimic on *IL-22RA1* expression, organoids were treated with 2ng/ml of recombinant ovine IL-22, BIO-RAD (PSP007) or murine IL-22 recombinant protein, PeproTech® (210-22-10UG). Organoids were transfected with miR-5352 mimic or miRNA control mimic at a final concentration of 50 nM for 1 hr at 37°C, after which the organoids were carefully mixed with Matrigel and transferred into appropriate culture wells with IntesiCult OGM media and corresponding inhibitors for ovine abomasum organoids or without inhibitors for murine SI tuft cell reporter organoids. Organoids were cultured for 2 or 4 days before RNA extraction, 4 days for immunofluorescence assay (IFA) and 12 days for phenotypic studies, in which IL-13 and miR-5352 mimic or control mimic were added every two days when media was changed. Images for phenotypic studies were taken every two days under EVOS M5000 Imaging System (Thermo Fisher Scientific) and inverted confocal microscopy.

### Immunofluorescence confocal microscopy

Immunofluorescence staining was carried out as previously described by O’Rourke et al.^90^ with some modifications. Briefly, abomasum and SI organoids were fixed in 10% paraformaldehyde for 20 min at room temperature. Following fixative removal and washing with IF buffer, permeabilization solution was added for 20 min. Primary antibodies (anti-POU2F3, anti-Ki67, anti-KLF4, anti-MUC2, anti-LGR5, anti**-**CTNNB1) all diluted in blocking solution (IF buffer containing 1% bovine serum albumin (BSA)), were incubated overnight in a humidified chamber at 4°C. Secondary antibodies goat anti-rabbit IgG (H+L) Cross-Adsorbed Secondary Antibody, Alexa Fluor™ 488 and 647 (Thermo Fisher) were incubated at room temperature for 1 h. ProLong Gold (Invitrogen) was added before viewing under a Confocal Zeiss LSM 880. Antibodies are detailed in **Supplementary Table S10**.

### RNA extraction and RT-qPCR

Total RNA was extracted using RNeasy Micro Kit (QIAGEN) from three wells of ∼ 300 organoids each, following manufacturer’s instructions. DNase I was added directly to the column membrane and incubated for 15 min at room temperature and RNA was eluted in 50 μl of RNase-free water. RNA concentration was determined using Qubit™ RNA High Sensitivity (HS) (Thermo Fisher). Samples were stored at −80°C until use. PolyA tailing of RNA and cDNA synthesis was carried out using the miRNA 1st-Strand cDNA Synthesis Kit (Agilent Technologies) according to the manufacturer’s instructions and SuperScript TM III (Invitrogen 18080044). RT-qPCR was performed using the Brilliant III Ultra-Fast SYBR QPCR Master Mix (Agilent Technologies 600882). Samples were analysed using the AriaMx-Real-Time machine and software (Agilent Technologies). All RT-qPCR reactions were carried out at least in triplicate from three biological replicates (organoids originating from three different animals) at day 2 after treatments. Organoids without IL-13 stimulation were included as control, and the expression of predicted miRNA target genes was standardized to expression of ovine Actin (*ACTB*) or murine Glyceraldehyde 3-phosphate dehydrogenase (*GAPDH*) as reference genes using the formula reported by Pfaffl et al.^91^. To find an appropriate reference gene for normalization of the qPCRs, four ovine and murine genes were analysed by two programs: BestKeeper^92^ and NormFinder^93^. The gene with Cp standard deviation ≤0.5 with coefficient of correlation ∼1 according to BestKeeper analysis and a low stability index according to NormFinder analysis was chosen (**Supplementary Figure S4**). Primer efficiency was calculated by standard curve using 1:10 serial dilution of cDNA samples. Statistical analyses and plots were generated using the fgStatistics^94^. PCR primer sequences are shown in **Supplementary Table S11**.

### Bioinformatic prediction of Hco-miRNA target genes

To identify potential target mRNAs of *H. contortus* secreted miRNAs, the 3’ UTRs of ovine (Oar_V4.0; NCBI GCF_000298735.2) and bovine (ARS-UCD 1.2; NCBI GCF_002263795.1) were obtained using the BioMart tool from ensembl.org^95^. Bovine 3’UTRs were included in target prediction as the bovine genome is currently better annotated than the ovine genome. MiRanda and RNAhybrid algorithms^59,60^ were used to predict target genes in both genomes for Hco-miR-5352 and the ten most abundant Hco-miRNAs secreted from *H. contortus* adult and L4 stage, a total of 23 Hco-miRNAs^34^. MiRanda parameters used were: i) strict seed pairing; ii) score threshold: 140; iii) energy threshold: −17 kcal/mol; iv) gap open penalty: −9; v) gap extend penalty: −4; vi) scaling parameter: 4. For RNAhybrid parameters used were: i) energy threshold −20 kcal/mol ii) no G:U in the seed iii) helix constraint 2 to 8. TargetScan (Version 8.0) was also used for supporting predictions, using default settings and canonical target sites of 7mer and 8mer^96^ predicted in bovine 3’UTR. Target genes predicted by at least two of the three algorithms were considered high confidence targets. The orthologs between ovine and bovine were obtained using eggNOG-mapper v2^97^. All the data was integrated using R/Bioconductor package ‘dplyr’ version 1.0.10^98^. Dr. Tom’s network platform of BGI (https://biosys.bgi.com) was applied to construct and visualize the biological-term classification and enrichment analysis.

### RNA-seq

Ovine abomasum organoids were exposed to four different conditions for 48 h before RNA extraction and sequencing i) medium alone (control) ii) Stimulated with IL-13 (IL-13) iii) Stimulated with IL-13 and transfected with miRNA mimic-control (control mimic+IL-13) iv) Stimulated with IL-13 and transfected with miR-5352 mimic (miR-5352 +IL-13). Four biological replicates were used for each condition (n=4). Total RNA was extracted using RNeasy Micro Kit (QIAGEN) from three wells of ∼ 300 organoids, following manufacturer’s instructions. The RNA concentration was determined using Qubit™ RNA High Sensitivity (HS) (Thermo Fisher), and quality and integrity determined using Bioanalyzer. For each sample, 1 µg of total RNA was used for RNA-seq analysis. All library synthesis and sequencing were conducted at BGI (Hong-Kong). DNBseq technology was used with a sequencing length of PE 150. RNA-seq data is available in Sequence Read Archive (SRA) accession number PRJNA1102187.

### Transcriptional analysis of abomasum organoids

Quality control of RNA sequencing data was performed on the raw reads using the filtering software SOAPnuke (version v1.5.2) with parameters -l 15 -q 0.2 -n 0.05^99^. The filtered clean reads were then aligned to the reference genome *Ovis_aries* (NCBI: GCF_000298735.2_Oar_v4.0). For this purpose, the hierarchical indexing for spliced alignment of transcripts (HISAT2, v2.0.4, http://www.ccb.jhu.edu/software/hisat) was used, based on Burrows-Wheeler transformation and Ferragina-Manzini (FM) with parameters --sensitive --no-discordant --no-mixed -I 1 -X 1000 -p 8 --rna-strandness RF^100^. Clean reads were aligned to the reference genome using Bowtie2^101^ (v2.2.5), using the following parameters-q --sensitive --dpad 0 --gbar 99999999 --mp 1,1 --np 1 --score-min L,0,-0.1 -p 16 -k 200, to allow gene or transcript expression to be quantified. RSEM^102^ (v1.2.8); parameters: -p 8 --forward-prob 0 --paired-end was used to calculate the gene expression level of each sample. The overall analysis of transcriptome sequencing was also performed, including the transcript randomness using a Perl script for sequence saturation analysis. After alignment, statistics of the mapping rate and the distribution of reads on the reference sequence were used to determine whether the alignment results passed QC. For principal component analysis (PCA) standardized method z-score (by row) and normalized read counts as method of grouping were used. Differential gene detection was studied by DESeq2 method^103^, based on the principle of negative binomial distribution. Volcano plots were generated using Log2FC>1 for IL-13/CONTROL and Log2FC>0.5 for MIMIC/CONTROL MIMIC conditions to identify genes altered by miRNA regulation, which can be a mild modulatory effect. According to the results of differential gene detection, the R package pheatmap (https://cran.r-project.org/web/packages/pheatmap/) was used. Reads were also mapped to the bovine genome (ARS-UCD 1.2); percentage mapping was lower than to the ovine genome. The R package clusterProfiler was used for GSEA analysis using GO:BP terms, and tuft or mucous cell gene sets^35^ with default settings.

## Acknowledgements

The authors gratefully acknowledge Colin Loney, Imaging Manager (Medical Research Council-University of Glasgow Centre for Virus Research) for help with imaging, Dr David Smith (MRI) for provision of ovine small intestinal organoids, and Dr Roz Laing (University of Glasgow) for provision of *Haemonchus contortus* adult parasites. We thank Prof Brian Shiels for comments on the manuscript. This work was supported by the Royal Society through a Newton International Fellowship (NIF\R1\201159) and McIntyre International Research Fellowship (University of Glasgow) to MGP, and by a Wellcome Collaborative Award (Ref 211814) to RMM, CB, ED, PJ, TNMcN.

## Author contributions

**MGP:** conceptualization, data curation, formal analysis, funding acquisition, investigation, methodology, validation, visualization, writing original draft preparation, review & editing. **VG:** methodology, resources, review & editing. **WA:** investigation. **FG:** resources, review & editing. **FH:** resources, review & editing. **TNMcN:** funding acquisition, resources, review & editing. **RMM:** funding acquisition, resources, review & editing. **PJ:** funding acquisition, resources, review & editing. **ED:** conceptualization, funding acquisition, supervision, review & editing. **CB:** conceptualization, funding acquisition, resources, supervision, writing original draft preparation, review & editing.

## Supplemental information

**Supplementary Table S1.** Ovine abomasal organoid RNA-seq results summary. RNA-seq output data, showing number of reads and mapping to reference genome *Ovis aries* v4.0.

**Supplementary Table S2.** Differential expression analysis (DESeq) of ovine abomasal organoids treated with IL-13 vs. untreated abomasal organoids (CONTROL) in culture for 48 h.

**Supplementary Table S3.** Differential expression analysis (DESeq) of ovine abomasal organoids transfected with Hco-miR-5352 mimic (MIMIC) vs. control mimic (CONTROL_MIMIC). Both were co-treated with IL-13 and cultured for 48 h.

**Supplementary Table S4.** Abomasal genes with opposite direction of regulation in ovine abomasal organoids treated with IL-13 versus no treatment, compared to IL-13 plus Hco-miR-5352 mimic versus IL-13 plus control mimic.

**Supplementary Table S5.** Gene set enrichment analysis (GSEA) for Biological Process domain (GO:BP) terms (ranked by NES value) of ovine abomasal organoids treated with IL-13 relative to control with no stimulation, after 48 h of culture. Relevant terms are highlight in grey.

**Supplementary Table S6.** Gene set enrichment analysis (GSEA) for Biological Process domain (GO:BP) terms (ranked by NES value) of ovine abomasal organoids transfected with miR-5352 mimic relative to miRNA control mimic transfection, both treated with IL-13 and cultured for 48h. Relevant terms are highlight in grey.

**Supplementary Table S7.** Bovine and ovine bioinformatic target prediction datasets for Hco-miR-5352, and expression from ovine abomasal RNA-seq.

**Supplementary Table S8.** Protein concentration of *H. contortus* ES different fractions.

**Supplementary Table S9.** *H. contortus* miR-5352 mimic and control sequences used in organoid transfection experiments.

**Supplementary Table S10.** Details of antibodies used in immunofluorescence experiments.

**Supplementary Table S11.** Primer sequences of genes of interest and reference genes for ovine and murine RT-qPCRs.

**Supplementary Figure S1.** Spheroid phenotype in GI organoids after *H. contortus* or *H. polygyrus* concentrated ES exposure for 12 h. Representative light microscopy images of ovine abomasal organoids, ovine small intestinal (SI) organoids, and murine SI organoids exposed to concentrated *H. polygyrus* (Hpoly) and or *H. contortus* (Hco) adult excretory-secretory (ES) products (10 µg/ml). Control shows organoids in growth medium alone and lower panels show organoids exposed to the flow-through of Hpoly or Hco ES after concentration (3kDa cut-off). White arrows indicate spheroid organoids identified by thinner membrane and more granular appearance. Scale bar = 300 µm.

**Supplementary Figure S2. (A)** Expression of predicted miR-5352 target gene *Klf4* in ovine abomasal organoids with no stimulation, treated with IL-13 alone, control mimic (miRIDIAN negative control sequence) + IL13 (alone, with Lipofectamine, or DharmaFECT reagent), or scrambled miR-5352 mimic + IL-13 (alone, with Lipofectamine, or DharmaFECT reagent). miRNA mimics at 100 nM final conc. **(B)** Expression of *Klf4* in ovine abomasal organoids with no stimulation, treated with IL-13 alone, control mimic (miRIDIAN negative control sequence) + IL-13 or scrambled miR-5352 mimic + IL-13, using DharmaFECT as transfection reagent and miRNA mimic at 25, 50, 100 nM. Log2 fold change of RT-qPCR values were compared with non-stimulated control in three independent biological replicates +/− SD. Statistical analysis was by ordinary one-way ANOVA with Tukey’s multiple comparisons test; **, P < 0.01. **(C)** Representative images of *Dclk1* tdTomato^+^ tuft cells in murine SI organoids, untreated (No stim), treatment with IL-13, control mimic (miRIDIAN negative control sequence) + IL-13, scrambled miR-5352 mimic + IL-13, and miR-5352 mimic + IL-13, using DharmaFECT as transfection reagent and mimics at 50 nM in OGM for 4 days at 37°C. **(D)** Number of *Dclk1* tdTomato^+^ cells in murine SI organoids at 4 days after treatment with IL-13, control mimic + IL-13, scrambled miR-5352 mimic + IL-13, and miR-5352 mimic + IL-13, using DharmaFECT as transfection reagent and mimics at 50 nM. Graph shows the average number of tdTomato^+^ cells from 5 images (250 organoids per treatment). Statistics calculated with a two-tailed t-test on mean of biological replicates (n=2) compared to untreated organoids. Scale bars: 100 µm.

**Supplementary Figure S3. (A)** Volcano plot of differential expression analysis (DESeq) in abomasal organoids treated with IL-13 plus control mimic vs. those treated with IL-13 alone and cultured for 48 h. x-axis represents the fold change of the difference after conversion to log2; y-axis represents the significance value after conversion to -log10. **(B)** Heat map showing expression level of genes from ovine abomasal organoids treated with IL-13 plus control mimic and IL-13 alone using normalised average read counts, standardized method: log (value+1). Colours indicate level of expression from low (blue) to high (red).

**Supplementary Figure S4.** Reference gene selection for RT-qPCR data normalization by BestKeeper and NormFinder. Ovine Actin (ACTB) and mouse Glyceraldehyde 3-phosphate dehydrogenase (GAPDH) were the genes that showed a cycle threshold (Ct) standard deviation ≤0.5 with a coefficient of correlation ∼1 according to BestKeeper analysis, and a low stability value according to NormFinder analysis (minimal expression variability).

